# A Theoretical Comparison of Alternative Male Mating Strategies in Cephalopods and Fishes

**DOI:** 10.1101/2022.06.03.494773

**Authors:** Joseph Landsittel, G. Bard Ermentrout, Klaus M. Stiefel

## Abstract

We used computer simulations of growth, mating and death of cephalopods and fishes to explore the effect of different life-history strategies on the relative prevalence of alternative male mating strategies. Specifically, we investigated the consequences of single or multiple matings per lifetime, mating strategy switching, cannibalism, resource stochasticity, and altruism towards relatives. We found that a combination of single (semelparous) matings, cannibalism and an absence of mating strategy changes in one lifetime led to a more strictly partitioned parameter space, with a reduced region where the two mating strategies co-exist in similar numbers.

Explicitly including Hamilton’s rule in simulations of the social system of a Cichlid led to an increase of dominant males, at the expense of both sneakers and dwarf males (“super-sneakers”). Our predictions provide general bounds on the viable ratios of alternative male mating strategies with different life-histories, and under possibly rapidly changing ecological situations.

## Introduction

Alternative male mating strategies have been observed in a wide range of animals, from birds, mammals, insects, fishes to cephalopods (Taborsky, 1994; 1997; 2001). In a number of fish species sneaker males have been observed (blennies: Ros et al., 2006; gobies: Drilling & Grober (2005); Marentette et al., 2009; wrasses: Alonzo & Warner, 2000; Stiver, 2015; Suzuki, 2008; cichlids: Ota & Kohda, 2006). These smaller males typically do not defend a harem, territory or nest as the dominant males of the same species do; instead, they sneak into the harem or territory defended by a dominant male and aim to achieve matings with females in a clandestine, “sneaky” manner (Fig. 1). Sneaker males are typically smaller than dominant males, and in some species colored and patterned like females, likely to make detection by dominant males more difficult. In several fish species, males can switch strategies, from sneaker to dominant, during the course of their lifetimes.

**Figure 1:**
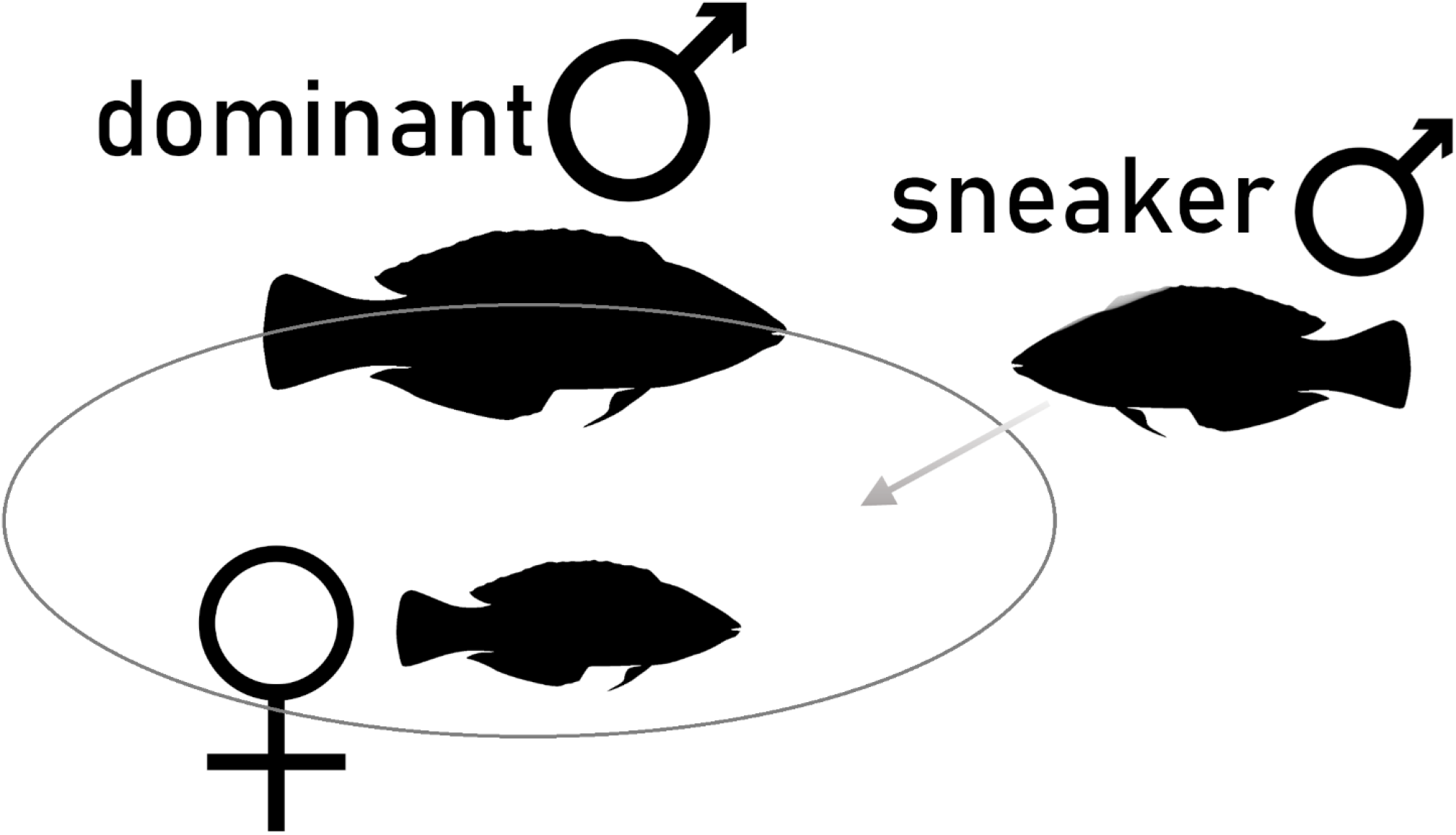
Schematic explanation of mating systems with dominant and sneaker males in fishes. A large dominant male defends a territory and/or one or more females. A sneaker male, to the right, attempts to gain access to the territory and/or females by surprising or deceiving the dominant male. Sneaker males are often smaller, and younger (earlier in their life-histories) than dominant males and can turn into dominant males in some species. Fish shape (the wrasse *Coris gaimard*) from Schiettekatte et al. (2019).

Interestingly, sneaker males also occur in phylogenetically completely unrelated cephalopods: cuttlefish, octopi and squid (Tsuchiya & Uzu, 1997; Norman et al., 1999; Hall & Hanlon, 2002; Ibáñez, 2019; Marian et al., 2019; Brown et al., 2021; Apostólico & Amoroso RodriguezMarian, 2020). In these species, just as in fishes, smaller males without a territory of their own clandestinely achieve matings with females, sometimes while taking up a female-like coloration (which is behaviorally plastic at fast time-scales in cephalopods). Since most cephalopods are semelparous (mate only once during a lifetime), strategy-switching typically does not occur in cephalopods; A number of other life-history strategy differences exist between cephalopods and fishes which likely influence their mating systems. These include the higher prevalence of cannibalism in squid, as well as generally very high growth rates in cephalopods.

We aimed to find a general explanation of the conditions which favor either the sneaker male or the dominant male strategy, as a function of the life histories of the animals in question, and as a function of global ecological factors (relative predation pressure,food availability and stochasticity). For this purpose, we used abstract mathematical models of the growth, reproduction and death of cephalopods, wrasses and cichlids. With these models, we determine the Evolutionary Stable Strategy (ESS), the ratio of sneakers to dominant males at equilibrium. For each of the three groups (wrasses, cichlids, cephalopods) that we simulated, we based our models on one particular species:

### Wrasses

The wrasses (Labridae; Froese & Pauly, 2021a) are a family of perciform marine fishes with a range of interesting reproductive behaviors, such as female to male sex change, territoriality and sneaker/dominant male strategies, with switches during a lifetime between these male mating strategies. We primarily based our model on the species *Symphodus ocellatus*, a Mediterranean wrasse with a well-studied mating system (Warner & Lejeune, 1985; Alonzo & Warner, 2000a, b).

### Cichlids

Cichlids are a family of mostly freshwater fishes with extensive adaptive radiations (Turner, 2007; Wagner et al., 2012; Brawand et al., 2014) in South America and the Great Rift Lakes of Africa. They show a wide variety of reproductive strategies, such as harems, mouth brooding, cooperative breeding and sneaker/dominant males (Taborsky, 2001). We specifically modeled the mating system of the African cichlid *Lamprologus callipterus*, which has a unique genetically determined class of sneaker males, which are about 40 times smaller in size than the dominant males. Due to a Y-chromosome (male inherited) marker, these dwarf sneaker males remain sneakers all their lives, in contrast to “regular” fish sneaker males which can switch to a dominant male role later in life (Schütz & Taborsky, 2000; Sato et al., 2004). We simulated the mating dynamics of this species due to the known, and unusual, role of genetic determinism in assigning males to either dominant/sneaker or dwarf sneaker male status.

### Cephalopods

In cephalopods, we primarily based our model on *Sepia apama*, the giant Australian cuttlefish, which has been thoroughly studied (Naud et al., 2004; 2005; Zylinski et al., 2011). Both dominant and sneaker males occur in this species which congregates yearly in South Australia for mass mating events (Hall & Hanlon, 2002).

### General Considerations

We initially based our abstract models of sneaker and dominant males on the known mating systems of the aforementioned species. While the models are initially based on a single species, due to the high level of abstraction, they nevertheless represent a range of species of the respective family or class with comparable life-histories. In the second part of the study, we hence compared the outcomes of the simulations to several wrasse and cephalopod species known to have sneaker and dominant males in their mating systems.

Fishes and cephalopods are phylogenetically only very distantly related, with their last ancestor being pre-Cambrian, and this ancient simple organism almost certainly was not capable of any sophisticated behavior akin to alternative mating strategies. Any similarities in the dynamics and trade-offs will not be due to a shared ancestry of the mating systems, rather due to common evolutionary pressures.

Using our models, we asked which conditions favor sneaker males, and which favor dominant males. Specifically:

1. How do food abundance and predation pressure influence the balance between sneaker and dominant males? How does stochasticity in food abundance influence optimal mating systems?
2. What is the consequence of semelparous mating (once per lifetime) in most cephalopods versus iteroparous mating (multiple times per lifetime) in many species of fishes? Notably, only iteroparous fishes can change between sneaker and dominant male strategies during a lifetime.
3. What is the consequence of cannibalism, known to play a significant role in schools of squid (O’Dor, 1998), but not in the fishes discussed here?
4. What is the consequence of including Hamilton’s rule (Hamilton, 1963; 1972) in our simulations, which states that an altruistic act is evolutionarily beneficial for an individual if it is sufficiently closely related to the receiver of the altruism?

## Methods

We used an abstract computational model to calculate the evolutionary stable strategies (ESS) regarding alternative male mating strategies that involve dominant and sneaker males. A mathematically detailed outline of our models is given in the appendix.

For the sake of simplicity, our model focused only on the dominant-sneaker male competition and did not include several other complexities of animal mating systems, even though they are highly important in the actual biological situations. The different variants of female choice during mating, especially during mating choices between dominant and sneaker males (Van den Berghe et al., 1998; Reichard et al., 2007) are not included in our model; In several species of wrasses, females seem to choose territories or nesting situations rather than individual males. In cephalopods, females can store sperm for a significant amount of time and seem to be able to regulate which sperm packages are used for fertilization (Sato, 2021). Also, the sperm morphologies between sneaker and dominant males are different in physiology and persistence in some species of fishes and squid (Hirohashi & Iwata, 2013). Lastly, in some species of wrasses, individuals can change from female to male after reaching a certain size threshold (Warner, 1982; Warner & Swearer, 1991). All of these complexities are of great biological importance and influence the mating successes of individuals. However, our aim is to focus solely on the effects of sneaker/dominant male mating strategies; including these additional complexities in our model would make it less tractable and clear.

Our models represent the fractions of the male population in several distinct states (Fig. 2): low energy, high energy, and mating for each of the dominant or sneaker variants. Death, as an irreversible state, is also explicitly included. The models alternate between a growth season, during which the animals can transition from a low to a high energy state, and a mating season. We ran the models until the populations reached equilibrium ratios of sneaker and dominant males, which we considered to be ESS. The models for wrasses, cichlids and cephalopods differ in the transition rates between states. Importantly, (1) there is no possible transition between a sneaker male and a dominant male state in cephalopods; (2) cephalopods are semelparous, meaning they mate once and then perish and therefore, the simulated cephalopod populations are terminated after one mating season. The third main difference between wrasses and cephalopods (3) is that the growth rate in cephalopods is twice that of the wrasses.

**Figure 2:**
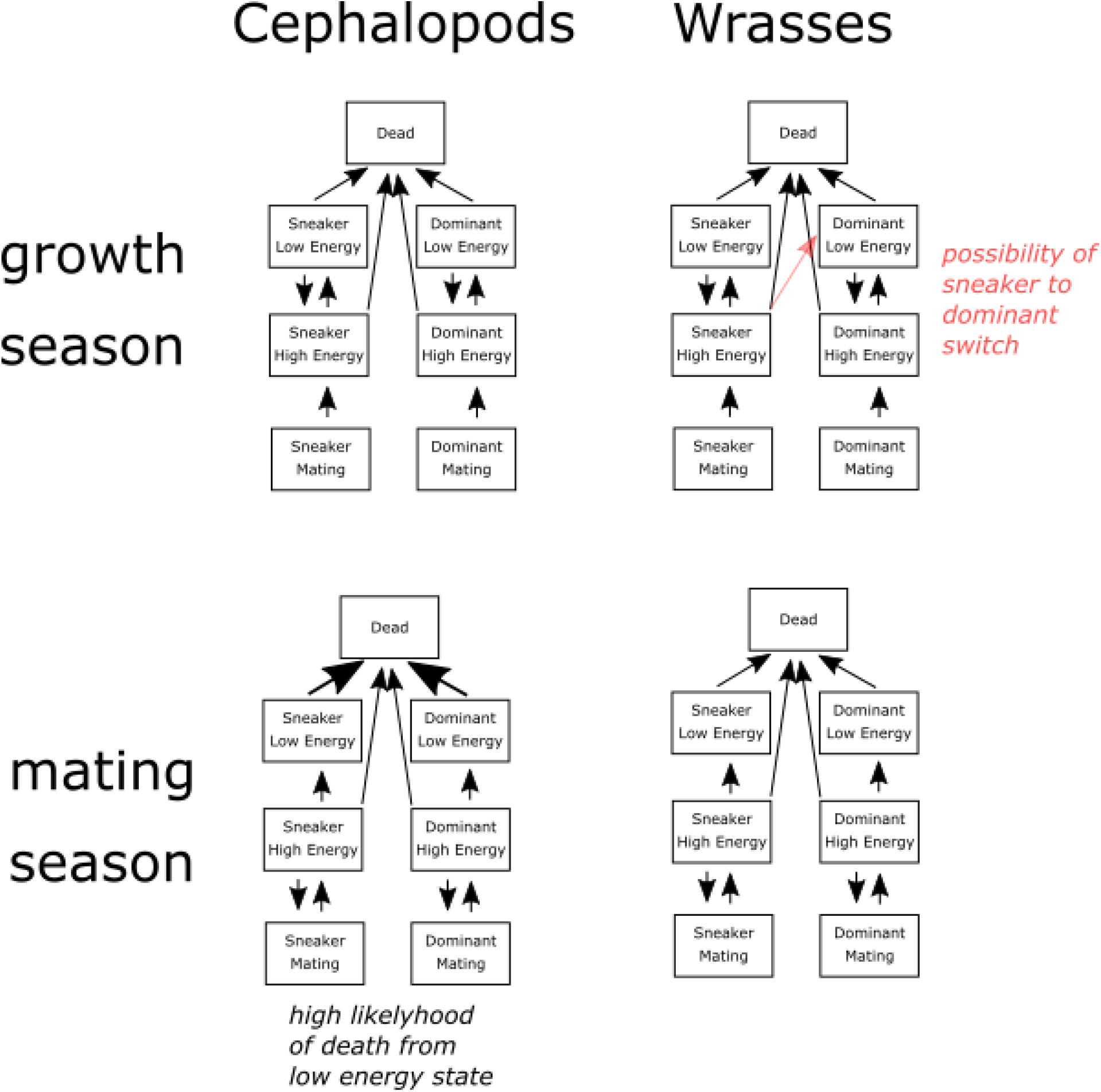
Flow diagrams outlining the transition between different states of the population of male animals in our models for cephalopods (left) and wrasses (right). The cichlid (*Lamprologus*) model contains a second column of dwarf sneaker males which can not switch to a dominant state (not shown).

The simulations representing the cichlid *Lamprologus callipterus* are identical to those of the wrasses but contain a second type of sneaker male (“dwarf males”), and hence a third set of states for these dwarf males exists. In addition, we explicitly included a bonus for dominant males which they receive from the success of – presumed closely related – dwarf sneaker males. Recall that these dwarf sneaker males are genetically determined (Y-chromosome linked) to be sneaker males of an unusually small size and are distinct from the “regular” sneaker males, which are not genetically distinct from dominant males and can change into dominant males later in life. The hypothesis that we aim to test with the simulations of *Lamprologus callipterus* is that the evolutionarily beneficial effects of providing altruistic acts – in this case providing nests – to close relatives contributes to the high incidence of dwarf sneaker males in this species.

Hamilton’s rule states that an altruistic act makes evolutionary sense if the donor and the receiver of the act are sufficiently related:

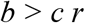

where *b* is the benefit of the altruistic act, *c* its cost and *r* the relatedness between donor and receiver. We assume in our simulations that *r* between dominant and dwarf sneaker males is high enough for this inequality to hold. Hence, we assume that the dominant males propagate their own genes by aiding the breeding of (maternally related) close relatives. We included this in the simulations by adding a bonus to the reproductive success of the dominant males proportional to the sneaker males when it came to calculating their proportion in the next generation.

Mathematically, our models are a *piecewise dynamical systems*, where a continuous development of variables alternates with (much rarer) breaks during which the dynamics change in a sudden manner (the mating/growth season transition). The parameters and mathematical details of our model are outlined in the appendix.

## Results

We first simulated the wrasse version of our model until it reached an evolutionary stable state (ESS), and performed a parameter sweep over different values of predation pressure (Fig. 3). We found that at very low predation pressures dominant males predominate; at intermediate predation pressures sneakers dominate; and at very high predation pressures dominants predominate again in the population. This result shows that the relative benefit of dominant or sneaker male reproductive strategy varies with ecological conditions in our simulations; this is in accord with field-observations which show that mating tactics are conditional on environmental factors (Engqvist, & Taborsky, 2006; Horth & Dodoson, 2007).

**Figure 3:**
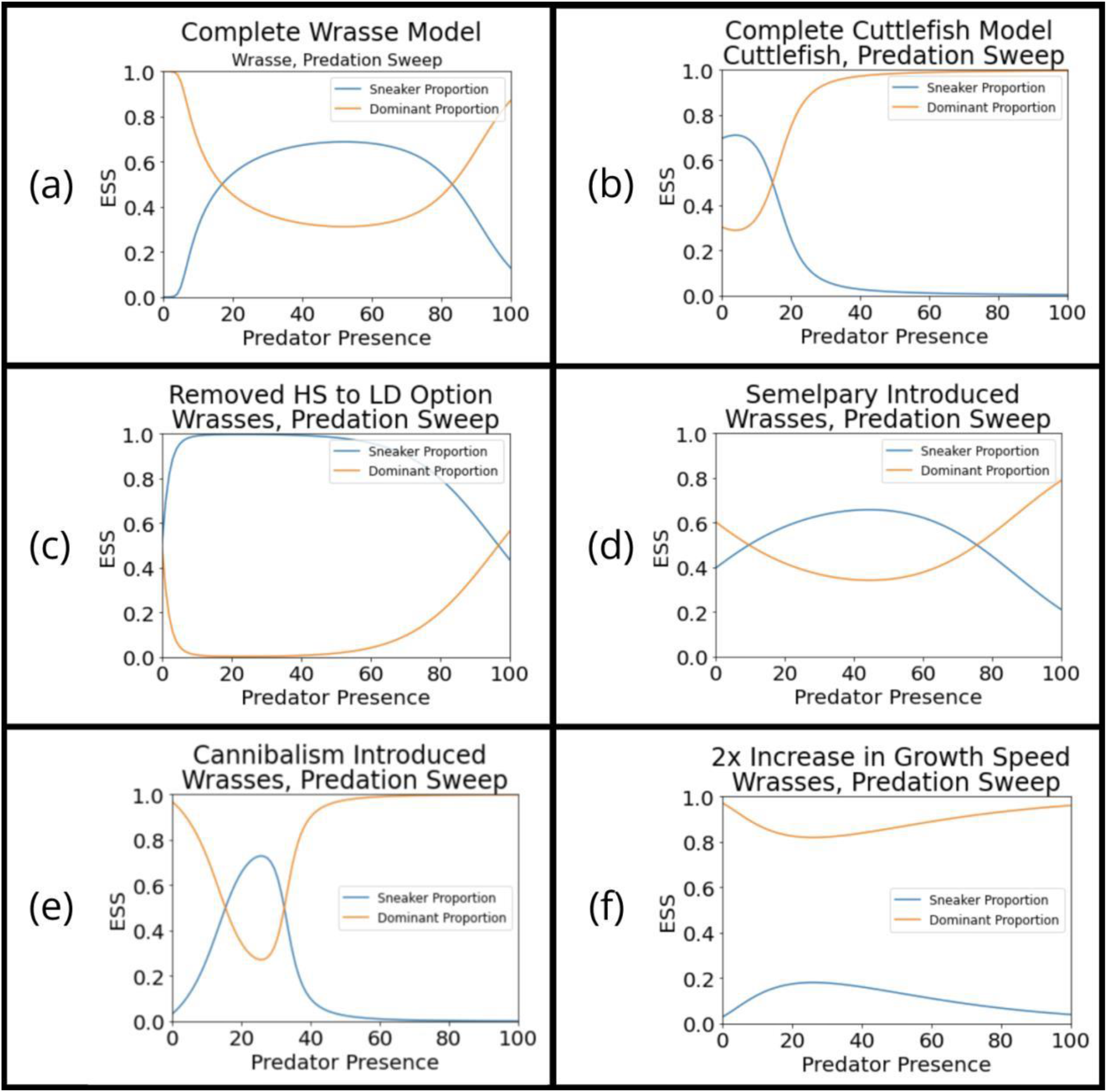
Changes from the model representing the wrasses to a model representing cuttlefish. a: Base wrasse model. B: Complete cuttlefish model. Each plot c-f shows the introduction of the individual differences (on their own, not sequentially) between the wrasse and the cephalopod model. Sweeps over varying amounts of predation pressure.

The differences between wrasses and cephalopods in our models are a faster growth rate, cannibalism, and semelpary in cephalopods. We then introduced these differences between wrasses and cephalopods individually into the model. Each difference was introduced on its own (not serially and cumulatively), to see what difference it makes on the relative proportion of dominant and sneaker males at an ESS (Fig. 3). We observed that removing the sneaker → dominant transitions eliminated the predominance of sneaker males at low predation pressures (Fig. 3b). Semelpary on its own flattened both the curve of the dominant and sneaker male proportions with increasing predation pressure (Fig. 3c). Adding cannibalism to the wrasse model flipped the curves: With this modification, dominant males, which are larger and more likely to cannibalize their smaller kin as opposed to vice versa, dominated the population at all but a narrow range of intermediate values (Fig. 3d). An increased growth speed compared to the base wrasse model moved the curves apart, and dominant males predominated at all predation values (Fig. 3e). The complete cuttlefish model introduces semelpary, cannibalism, the lack of sneaker to dominant transition and a higher growth rate all at once. It looks significantly different from the wrasse model: sneaker males predominate at low predation pressures, and dominant males at all other predation pressures (Fig. 3f).

The curves describing the relative proportions of sneaker and dominant males are almost flipped between the wrasse- and the cephalopod simulations. In the case of the wrasses, they are inverted u-shaped and u-shaped, and overlap at intermediate predation pressures; hence dominant males are in the majority at low and very high predation pressure. In the case of the cephalopods the curves resemble a sigmoid and an inverted sigmoid. The curves intersect in the center, hence sneaker males predominate at low predation pressures, dominant males at intermediate to high predation pressures (Fig. 3).

Our models predict that different ecological situations (predation pressure) will favor different proportions of sneaker and dominant males, which is in agreement with the empirical literature; our models also predict that the change in sneaker to dominant male ratio will be very different for wrasses and cephalopods.

### Two-parameter sweeps of Predation Pressure and Food Availability

The parameter sweeps described in the previous section were run at intermediate values of food availability. We also compared two-parameter sweeps between the model of the wrasses and the cephalopods, varying both predation pressure and food availability (Fig. 4). We observed that the region with a high proportion of sneaker males (in white) follows an inverted u-shaped outline in the case of the wrasses (Fig. 4a), indicating that over a wide range of food availabilities sneaker males dominate at intermediate predation values. In contrast, in the case of the cephalopods the region with a large proportion of sneaker males is found in the bottom right corner of the plot, with low predation pressure and low food availability. These plots confirm that the differences in life-histories between wrasses and cephalopods lead to a significant difference in which alternative male mating strategy is preferred at a given ecological situation.

**Figure 4:**
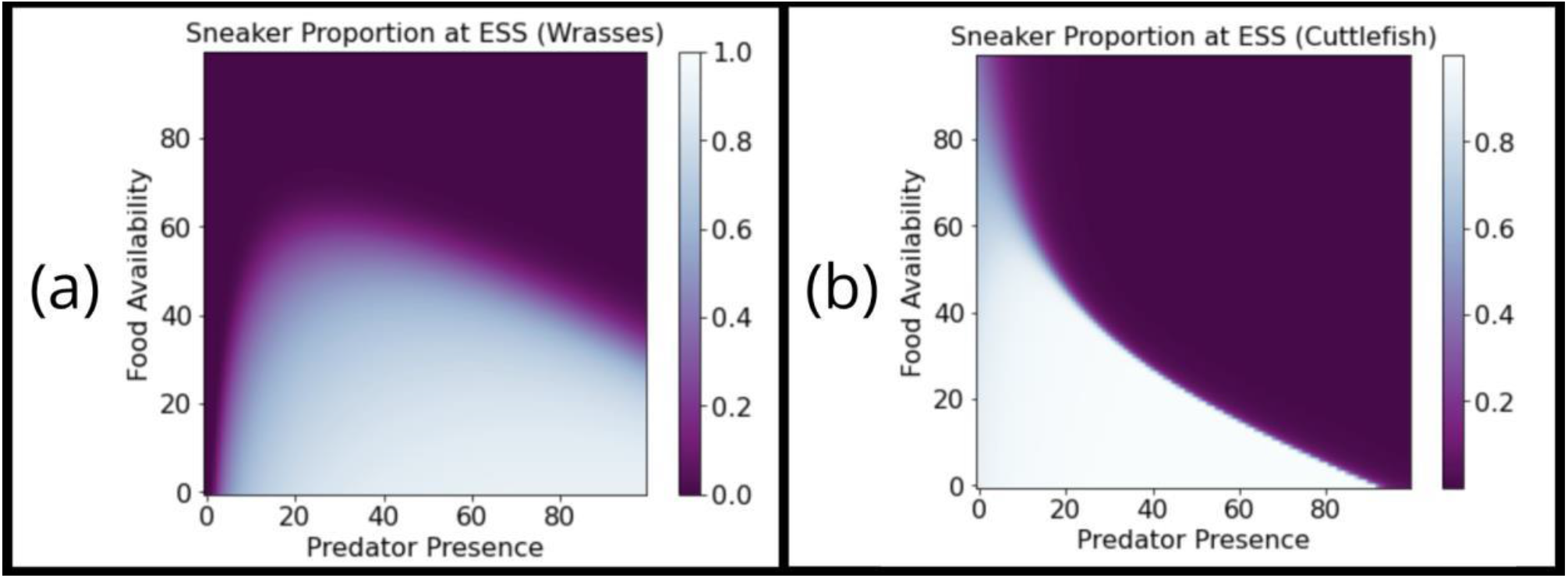
Two-parameter sweeps showing the proportion of dominant males (blue) and sneaker males (white) at ESS as a function of food availability and predation pressure for wrasses (a) and cephalopods (b). Each point represents one simulation run until it reached an evolutionarily steady state (ESS).

### Stochasticity

In real-world ecosystems, food availability is often stochastic, with the amount of food varying over time. The level of stochasticity varies greatly between different ecosystems. The evolution of life-history parameters can buffer the effect of environmental uncertainty (Wilbur & Rudolf, 2006). We hence tested how the ESS changes as a function of increased stochasticity of food availability. The transition probability (see appendix) from low-to high energy state was randomly changed between the time-steps of the simulation.

We found that increased stochasticity of food availability increases the proportion of sneaker males (Fig. 5); random events enable both sneaker males to persist at higher food availabilities and dominant males at lower food availabilities; but the effect is asymmetrical: The increase of sneaker males is more pronounced than the increase of dominant males (Fig. 6). This is due to a “bottoming out” effect, with the increase in dominant males happening in an unfavorable part of the parameter space where food availability is low.

**Figure 5:**
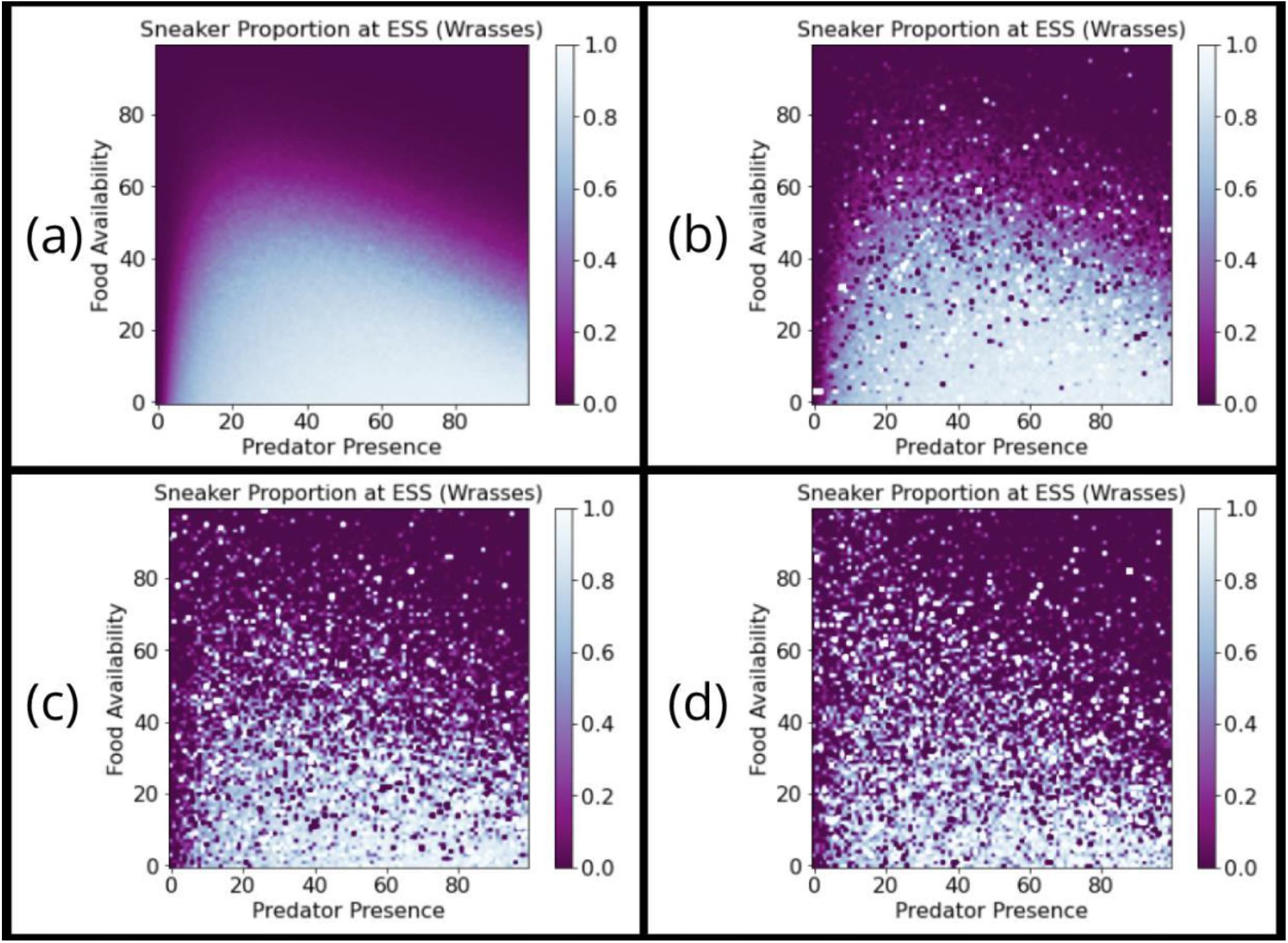
ESS with increasingly stochastic of food availability. Two-parameter sweeps showing the proportion of dominant males (blue) and sneaker males (white) at ESS as a function of food availability and predation pressure for wrasses. Each point represents one simulation run until it reached an evolutionarily steady state with stochastic food availability (ESS). Plots a to d illustrate increasing levels of stochasticity.

**Figure 6:**
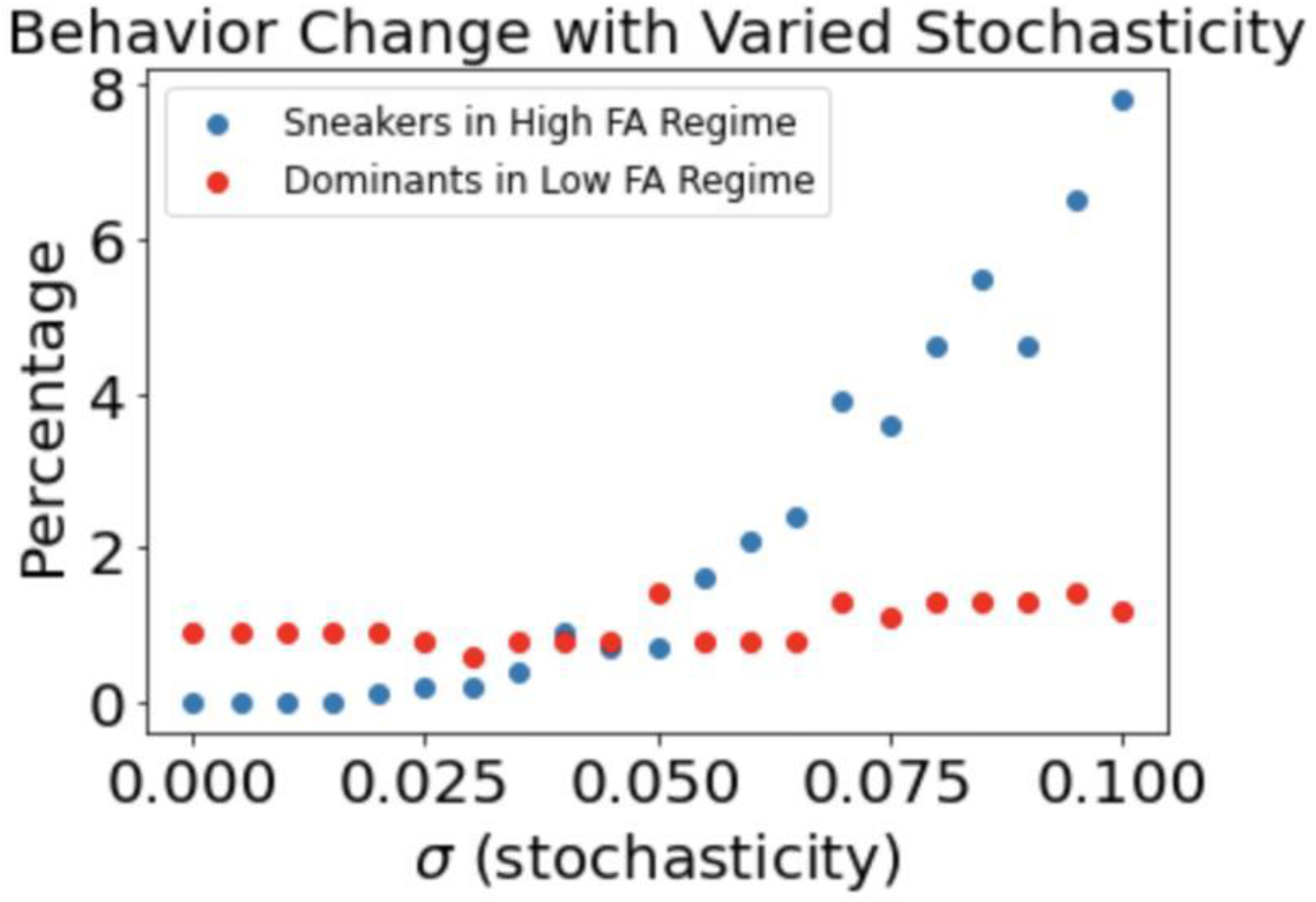
ESS of sneaker males and dominant males of wrasses at increasingly stochastic food availability, at intermediate predation pressure.

### A Cichlid with Genetically Determined Sneaker Males

Finally, we investigated how adding a genetic component to the mating system will influence the relative ratio of dominant and sneaker males. In the aforementioned cichlid *Lamprologus callipterus* a distinct group of dwarf sneaker males exist which are up to 40 times smaller than both the dominant and regular sneaker males and these remain sneaker males all their lives due to a genetic marker inherited in the male line. We hypothesized the high proportion of dwarf sneaker males in this species is partially due to their close relatedness with the dominant males. Hence, a dominant male which builds a nest that is then used to rear the offspring of a dwarf sneaker male will contribute to the rearing of offspring he is related to; genetically the effort made in building the nest is not lost.

We simulated the effect of the genetic relatedness by adding a bonus to the dominant male reproductive rate derived from Hamilton’s rule describing the beneficial effects of altruism (Fig. 7). This bonus represents the indirect reproductive success by facilitating the reproduction of relatives (dwarf males). Unsurprisingly, a bonus conveyed to dominant males increased their proportion in the population. This was especially the case at intermediate food availabilities and predator presences, and at the expense of both regular sneaker males as well as dwarf sneaker males.

**Figure 7:**
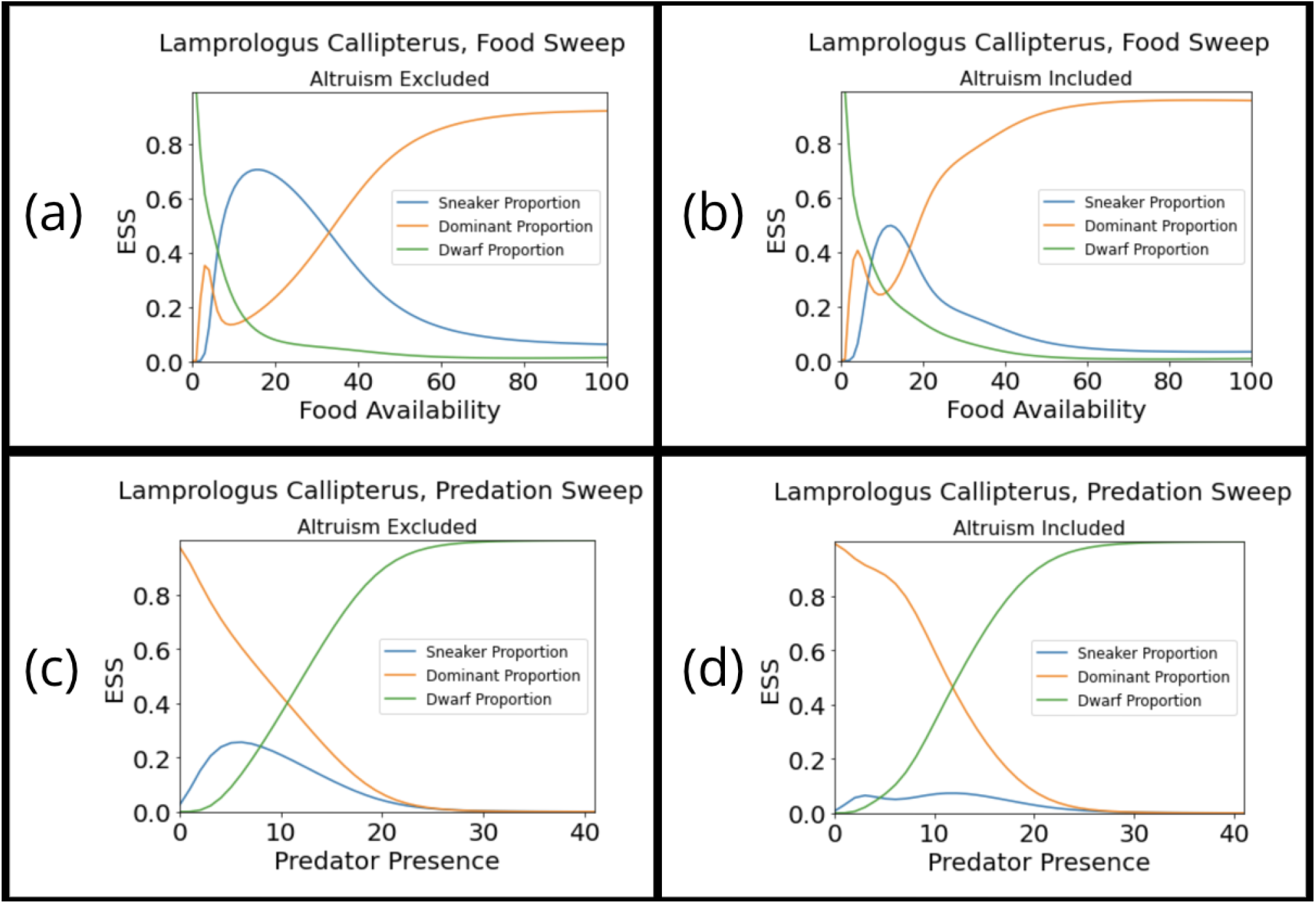
The effect of explicitly rewarding altruistic acts from dominant males to dwarf sneaker males (b, d) versus control simulations without the inclusion of altruism (a, c). Simulation parameters were varied over food availability (a, b) and predator presence (c, d).

## Discussion

We found that semelpary, cannibalism, a lack of switching between sneaker and dominant strategies and increased growth speeds, as seen in cephalopods compared to fishes changes the relative proportion of sneaker males as a function of ecological parameters (Fig. 8). At low predation pressure, sneaker males predominate in cephalopods, while dominant males predominate in wrasses. Two-parameter sweep over predation pressure and food availability confirm the fundamentally difference between the optimal sneaker:dominant male ratios for different ecological conditions. We also found that increasingly stochastic food availability favors sneaker males.

**Figure 8:**
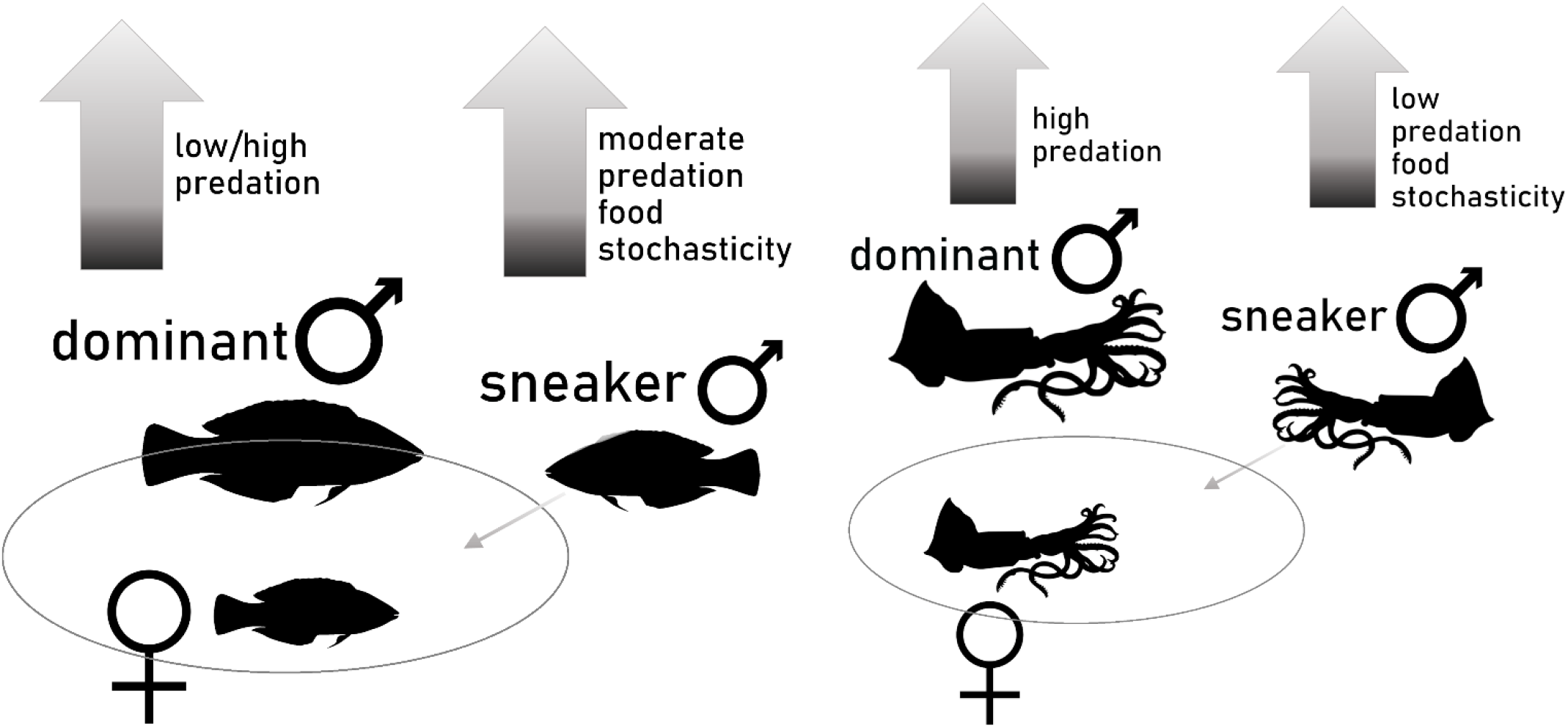
Schematic explanation of the results of our simulation study. In wrasses (left) dominant males are favored at low and very high predation levels. Sneaker males are favored at moderate predation levels and when food availability is stochastic. In cephalopods (right) dominant males are favored at high predation pressures, and sneaker males are favored at low predation pressures and when food availability is stochastic.

Finally, we found that an explicit inclusion of the genetic benefits of altruistic acts as described by Hamilton’s rule in the simulated mating system of the cichlid *Lamprologus callipterus* increases the proportion of dominant males, at the cost of both sneaker and dwarf sneaker males. Previous theoretical studies have already suggested that altruism can be a driver of the evolution of life-histories and social systems (Stiefel, 2013, 2014; Bourke, 2014).

The question is how these theoretical results can be empirically verified. Controlled experiments with populations of often widely migrating animals at evolutionary time-scales are impossible. We can, however, plot known proportions of sneaker to dominant males in mating systems on the 2-dimensional parameter sweeps with we conducted (Tab., 1, Fig. 9). When plotting the observed sneaker:dominant male ratios for wrasses (Fig. 9a) and cephalopods (Fig. 9b) each value becomes an isocline on the 2-parameter sweep plots: Each ratio is equivalent to an altitude on the surface describing the outcomes of the simulations, hence the simulations predict each ratio for a set of predation pressure and food availability values. As the field observations of sneaker:dominant male ratios for both cephalopods and wrasses vary widely, the isoclines are located in vastly different regions of the 2-dimensional parameter sweeps.

**Table 1:** Examples of cephalopod, labrid and cichlid species with sneaker:dominant male ratios observed in the field.

| Species | male:female | dominant:sneaker | Location | reference | Comment |
| --- | --- | --- | --- | --- | --- |
| **Cephalopoda** |  |  |  |  |  |
| *Sepia apama* | 11:1 to 4:1 | 1 : 5 | Whyalla, South Australia, 33 deg south | Naud et al., 2004 |  |
| *Loligo vulgaris* | 1.4:1 | 6 : 1 | Port Alfred, South Africa, 33 deg south | Hanlon et al., 2002 |  |
| **Labridae** |  |  |  |  |  |
| *Symphodus ocellatus* | 1.96 : 1 | 2 : 1 | Corsica, France, 41 deg north | Alonso, 2000 | satellite and non-breeding males not counted |
| *Thalassoma bifasciatum* | 1:2.7 | 1:1 | US Virgin Island, 18 deg north | Warner & Swearer 1991 | immature males assumed to be sneakers |
| *Thalassoma lucanasum and bifasciatum* | 1:1 to 1:10 | 1:8 to 1:99 | Panama, 9 deg north | Warner & Hoffman 1980 | sneaker ratio depending on population size |
| *Thalassoma lucasanum* | 1:1 | 1:5.8 | Panama, 9 deg north | Warner, 1982 |  |
| **Chiclidae** |  |  |  |  |  |
| *Lamprologus callipterus* |  | 1 : 2.27 | Lake Tanganyika, Tanzania, 6 deg south | Ocana, 2014 | freshwater; "dwarf male sneaker" ratio given |

**Figure 9:**
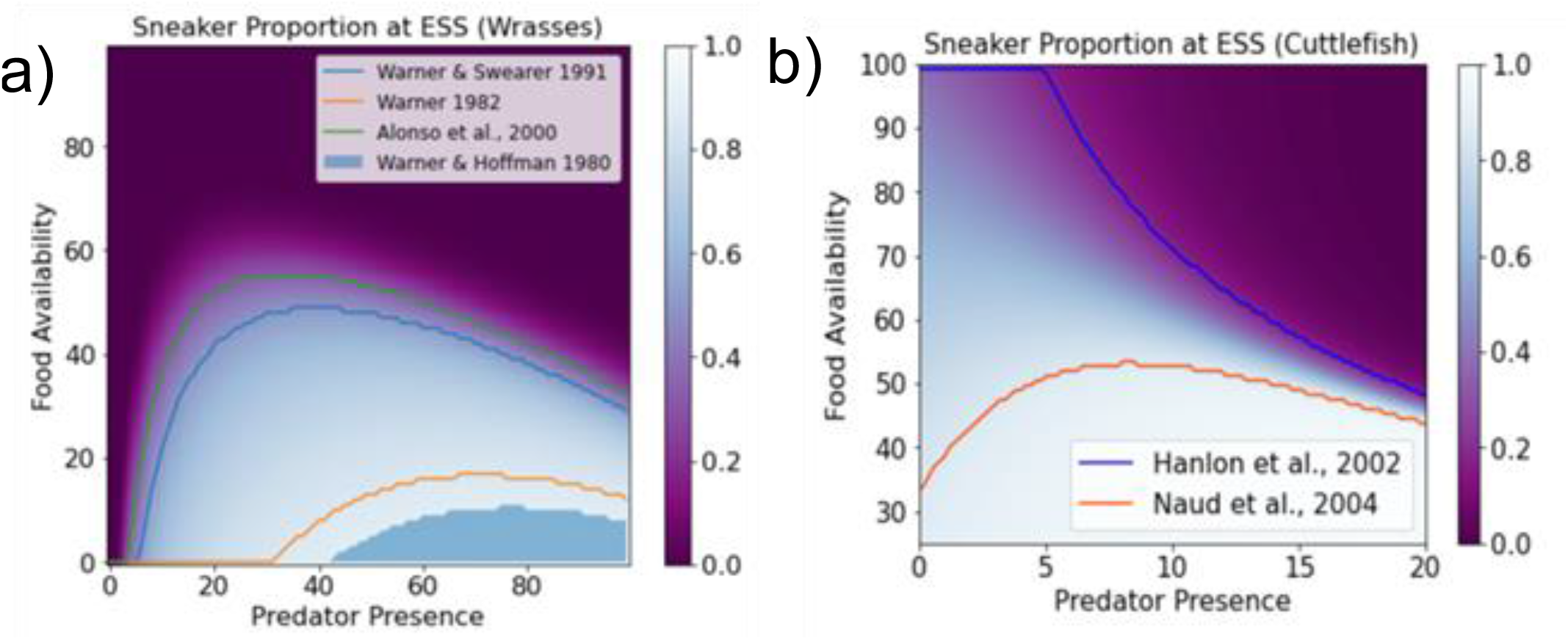
Proportions of dominant and sneaker males onto the two-parameter sweeps from our simulation results compared to observations from field studies of wrasses (left) and cuttlefish (right) published in the literature. Each isobath corresponds to one observed proportion in one species.

While the units of our predation pressure axis are necessarily arbitrary, they allow to assess the effect of *relative* changes. A decrease of predation pressure from moderate/high to low should lead to an increase of sneaker males in cephalopods, but to an increase of dominant males in wrasses. Predation pressure can in- or decrease due to a variety of reasons, such ecological changes along the geographic range of a species, or due to human intervention. Generally, predation is increased in tropical/warmer and shallower oceans (Ashton et al., 2022). Human over-fishing can re-organize marine ecosystems, often decreasing the number of top-level predators and conversely increasing the number of mid-sized predators (“meso-predator release”, Ritchie & Johnson, 2009). These change in predation pressure experienced by a species with alternative male mating should lead to changes in the sneaker to dominant male ratio as predicted by our simulations.

Mating systems are subject to evolutionary pressure like other organismal traits such as morphological or physiological features of an animal (Taborsky, 2001). Evolutionary responses of mating systems to environmental variables such as resource variation can be complex; however, the responses of mating systems to a combination of environmental and life-histories will inevitably be more complex. The use of numerical simulations in this study hopes to elucidate these complex dependencies, in comprehensive ways which are impossible via field observations or laboratory experiments.

## Figures

**Figure Appendix 1:**
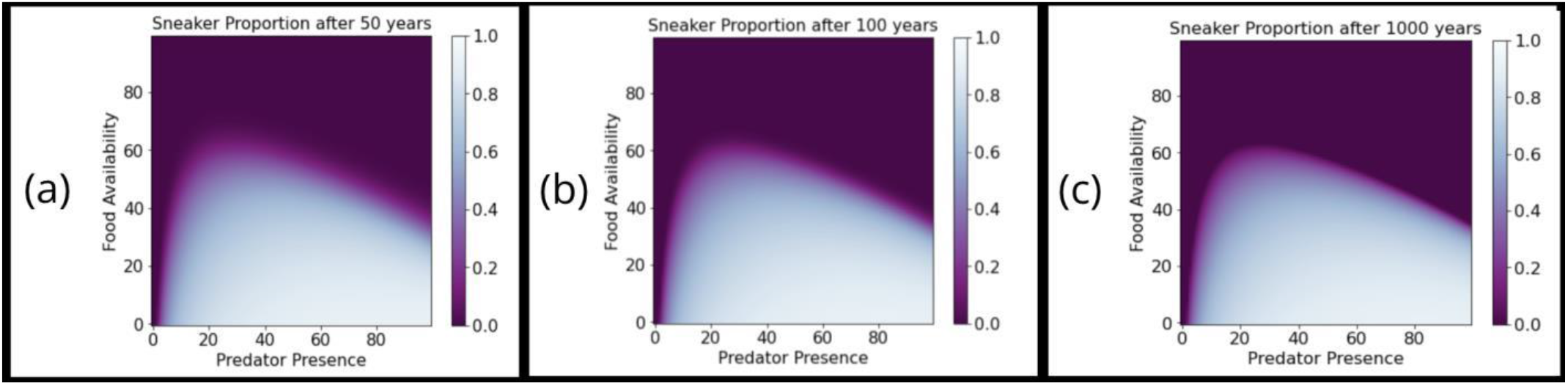
Comparison of the state of the 2-dimensional parameter sweeps after 10, 50 and 1000 (a, b, c) simulated years.

## Appendix: Equations and Parameters of the Models

Typically, models simulating the evolution of populations of animals are based on *dynamical systems*. In such a dynamical system a state vector, ***x***, represent the fractions of a population. If the population is divided into *n* unique states, we can view ***x*** as lying in a phase space *x* ∈ *X* ⊂ *R*^*n*^, and the population can be seen as evolving according to a map, *φ*:

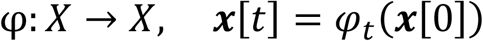

Such a mapping can either be discrete (*i*.*e*., *t* ∈ ℤ) or continuous (*i*.*e*., *t* ∈ ℝ). Additionally, it may or may not be smooth over time. In other words, it is conceivable that the population will evolve in different ways depending on separate periods or “seasons”. In our model, we simulate a mating season alternating with a growth season. The evolution of the state during the growth season will depend on how the state evolved during the previous mating season.

The model we used is a *discrete, piecewise-smooth dynamical systems*. Each timestep represents one day, and at each step the population is updated according to a time dependent mapping. With µas a set of parameters, the master equation of our model is:

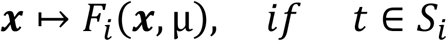

Separating the growth and mating season in our model is implemented as:

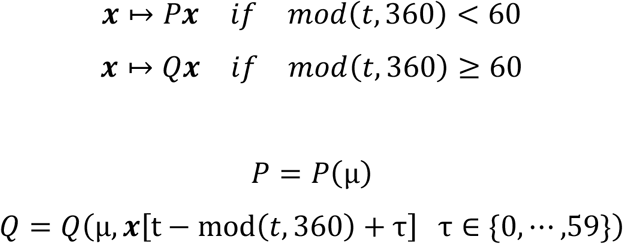

Where *P* and *Q* are matrices, each depending on a set of parameters, µ. *Q* also depends on the values taken on by the state vector, ***x***, during the previous mating season. The evolution of a state vector according to a transition matrix such as this is generally referred to as a *Markov chain*.

Figures including results iterate this by applying *P* 60 times and then *Q* 300 times for a total of 50 cycles. Figure A1 demonstrates that 50 years is sufficient for producing the same behavior as is present in 100 year and 1000 year simulations (Fig. appendix 1).

### Base Model: Wrasses

To model the alternative male mating strategies for wrasses, we used a piecewise system of Markov chains (Fig. 1). On any given day, a fish can move from state x to state y with probability P_x→y_. A one by seven state vector ***x***[t] is initialized as follows:

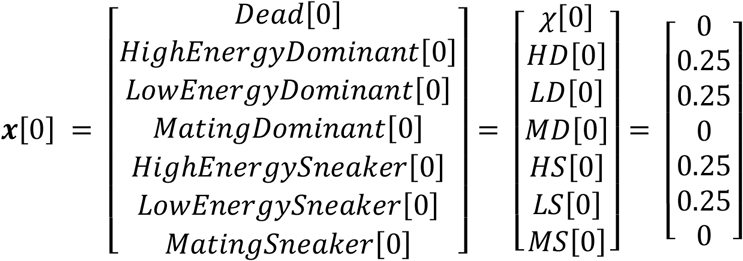

The population at t = 0 is assumed to be uniformly distributed across the high and low energy states for the two mating strategies (dominant/sneaker). The population is simulated with two transition matrices: *P* is applied to ***x***[t] for the first 60 days (during the mating season), and then *Q* is applied to ***x***[t] for the following 300 days (during the growth season).

The values of ***x***[*t*][*i*] take on real numbers between zero and one representing the proportion of the population that any given state represents. Hence the following holds true for all *t*:

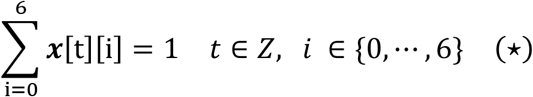

During the mating season:

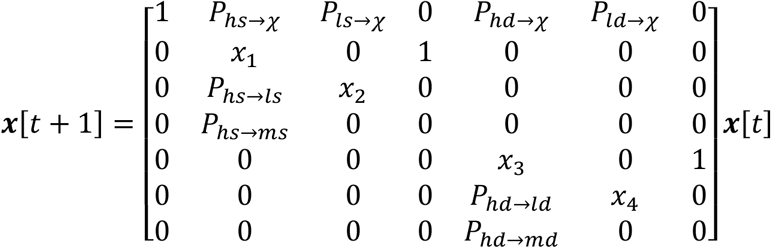

During the growth season:

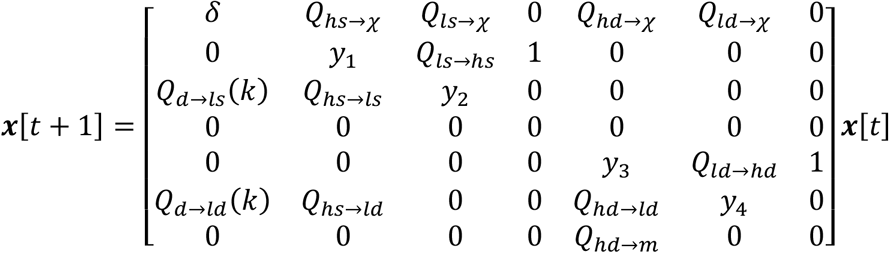

The element in the location *(i,j)* of a transition matrix is the probability that a subject in state *i* will transition towards state *j* in one day. The *i*^*th*^ state is defined as the state corresponding to the *i*^*th*^ entry of ***x***[t]. Observe that the placement of zeros is consistent with the diagram given by figure 1. Making choices for these parameter values requires a consideration of both (1) how the previous mating season played out and (2) to what extent predation and food are readily present in the environment.

To incorporate (1) into the model, we utilize *Q*_*d*→*ls*_ and *Q*_*d*→*ld*_. More accurately, we devised a deterministic method for pulling subsets of the population out of the dead state and into the two low energy states. This re-drawing of individuals from a pool of the dead is a mathematical approach reflects the birth of new fish during the growth season resulting from mating in previous months, devoid of biological interpretation. We used the following method for choosing *Q*_*d*→*ls*_ and *Q*_*d*→*ld*_ at the beginning of the growth season for the *k*^*th*^ year of iteration:

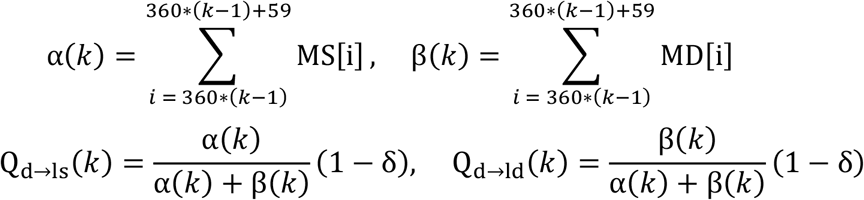

The likelihood that a new sneaker is born depends explicitly on the relative fraction of matings by sneakers in the previous cycle relative to dominant males. δ ∈ (0,1) reflects volatility in the population, or the proportion of the dead pool distributed towards sneakers and dominants during the subsequent growth season. We choose *δ* = 0.2.

The remaining transition probabilities were chosen according to the following relationships:

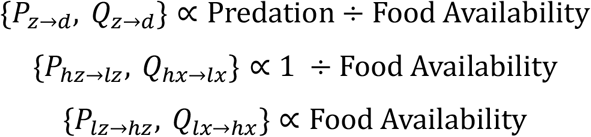

Here z corresponds to an arbitrary state. More specifically, let *α* correspond to food availability and *β* predation. The transition probabilities are as follows:

*Low Energy Sneaker Transitions:*

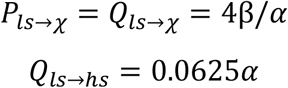

*High Energy Sneaker Transitions:*

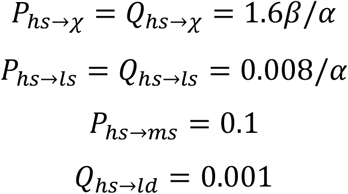

Low Energy Dominant Transitions:

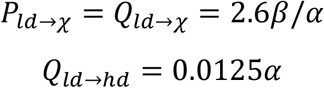

*High Energy Dominant Transitions:*

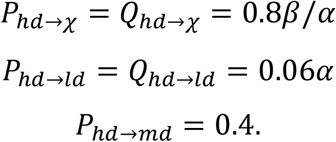

For the two-dimensional parameter sweeps (figures 4 and 5), *α* was swept from 1.20 to 2.00, *β* from 10^−5^ to 0.02. This choice magnifies the curve of change in ESS. These have been re-scaled to a 1 to 100 scale for the purpose of figures.

The specification of these proportionality constants was done with the following principles in mind:

- The sneaker male is more likely to increase its energy level in a given day than the dominant male.
- Though the dominant male is slower to evolve, it is less likely to subsequently loose energy.
- The sneaker male is more likely to die than the dominant.
- The sneaker male is less likely to transition from the high energy state to the mating state than the dominant male.

Continuing to fill in our matrix, we must consider the values along the diagonals, which describe the probabilities that a subject does not change its state. There is no immediately obvious biological intuition for these values. The columns of each transition matrix need to sum to one. Hence, we will choose each *x* and *y* such that this property is necessarily true.

The last parameter to determine is *Q*_*hs*→*ld*_: the frequency for which a male mating strategy change occurs. For the purpose of this model, we take this as a constant property for any in the model describing the of wrasses. The mating strategy change is known to depend on environmental variables in some species, however this dependence is not included in our model.

In the simulations the updates of the values of the matrix were iterated for 50 years. Note that a simulated year does not correspond directly to a chronological year, since the simulations were set up to reach an equilibrium at the fastest possible rate and aim to reproduce the equilibrium, but not the path to equilibrium of biological evolution. In Fig. A1 we compare the state of the simulations after 10, 50 and 1000 simulated years, and observe that the sharpness of the transition between sneakers and dominant males, but not the qualitative shape of the result changes.

### Cephalopods

We model cuttlefish differently from wrasses by instituting four characteristic changes. First, the sneaker to dominant transition is removed as cuttlefish are not observed to change strategies. Next, cuttlefish are observed to be semelparous, so the state of the population undergoes a mass death directly following every mating season. This is executed by resetting the state vector, *X*[t], to be heavily biased towards the dead state once per year. The model does not reset though, given that the information from the previous mating cycle will still be encoded in the parameters *Q*_*d*→*ls*_ and *Q*_*d*→*ld*_. Thirdly, a cannibalism mechanism is introduced. This is executed by letting *Q*_*ls*→*d*_ and *Q*_*hs*→*d*_ be dependent on the total population of cuttlefish playing the dominant strategy at any given time. More specifically, at each time step, *t*, these two transition probabilities are redefined in the following way:

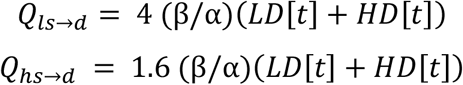

Additionally, the values of *y*_1_ and *y*_2_ are updated appropriately at each time so that (⋆) is not disrespected. Finally, the growth rates of cuttlefish are higher than those of fishes. Subsequently, we increased the values of *Q*_*ls*→*hs*_ and *Q*_*ld*→*hd*_ by a factor of two.

Figure 3 includes parameter sweeps over varied predation that result from each of these four changes applied individually to the basal wrasse model. The normalized food availability of *α* = 1.52.

### Lamprologus callipterus

Lastly, we shall consider the case of *Lamprologus callipterus*. A third mating strategy, the dwarf male, is introduced as a genetically determined sneaker. We do this by introducing three new dimensions for our state vector and transition matrices:

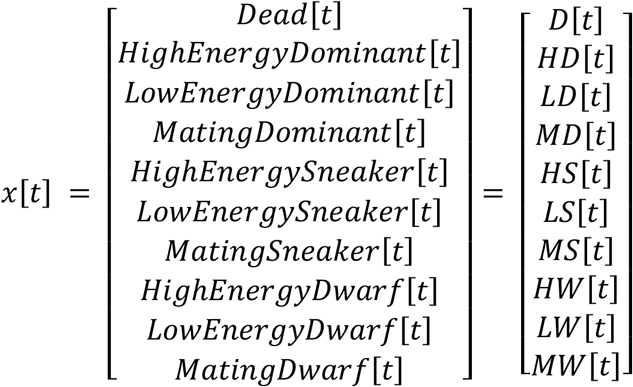

And during the mating season:

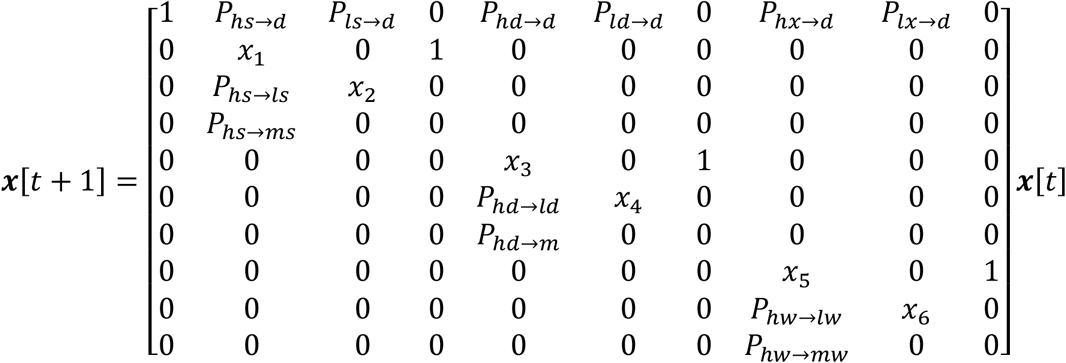

During the growth season:

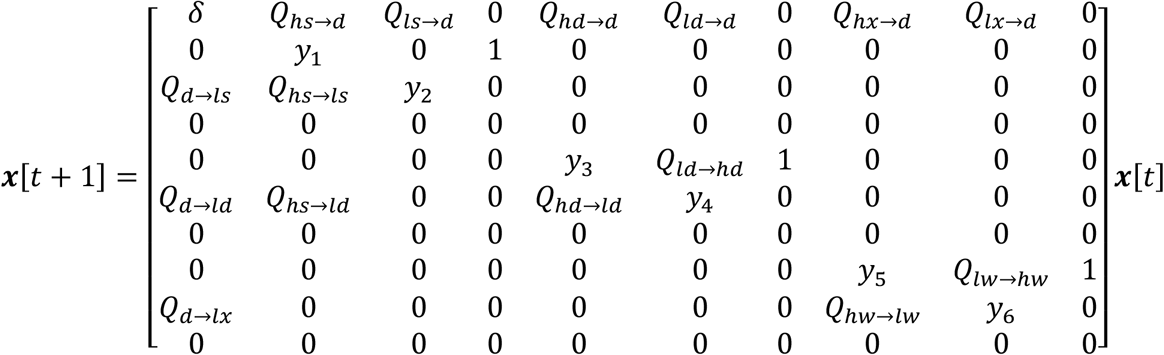

We also included a link between the dominant and dwarf strategies. Specifically, the dwarf males are assumed to be genetically related to the dominant males. This leads to an incentive for altruistic behavior for the dominant males.

To incorporate this, we introduced Hamilton’s rule by assuming a mean relatedness between any two male members of the breeding population. The dominant *Lamprologus* individuals receive a benefit for their altruism that is proportional to the total number of dwarf male in the population. This benefit will materialize as an increase to their probability of transitioning to the high energy state. At every time interval we performed the following update:

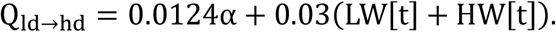

The remaining transition probabilities are given as follows

Low Energy Dwarf Transitions:

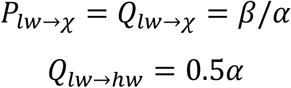

*High Energy Dwarf Transitions:*

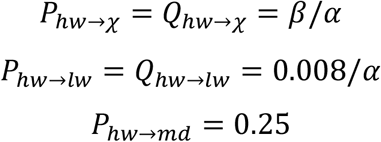

Given these values, one could fill in the transition matrices *P* and *Q* and simulate the population. Figure 7 includes the resulting ESS incorporating parameter sweeps over predator presence and food availability (with and without the altruism bonus). In 7a and 7b, predation is fixed at *β* = 0.001; in 7c and 7d, food availability is fixed at *α* = 1.0. In both cases, the other parameter was swept over the same aforementioned ranged as for the wrasses.

### Stochastic Considerations

We recognize that food availability and predator presence are not strictly fixed parameters in practice, but instead quantities that fluctuate with time around some initialized state. This is materialized in our model but introducing stochasticity in the low to high and high to low energy transitions.

Consider the baseline wrasse model. We can introduce multiplicative noise with a stochastic variable, *ξ*, which acts as a multiplier on the low to high energy transitions and a divider on the high to low transitions. *ξ* > 1 corresponds to a relatively high concentration of food and relatively low predator presence, specifically due to environmental variability. *ξ* < 1 is interpreted as the natural inverse. *ξ* is chosen via the following mechanism: at the beginning of a new season (mating or growth), reset *ξ* = 1. For each following day, update the variable according to the following equation:

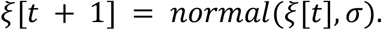

That is, a new value of *ξ* is chosen from a random normal distribution with mean *ξ* and variance *σ*^2^. In other words, *ξ* takes a random walk during each season before being reset. During a mating season, transition probabilities are updated in the following way each day:

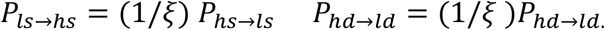

Similarly, during the growth season:

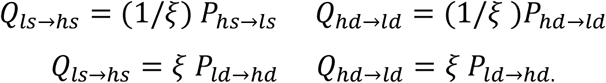

We can then run simulations to observe that the sneaker or dominant strategy may be present in parameter regimes despite being unstable in the deterministic sense.

